# Hyperexcitable phenotypes in iPSC-derived neurons from patients with 15q11-q13 duplication syndrome, a genetic form of autism

**DOI:** 10.1101/286336

**Authors:** James J. Fink, Jeremy D. Schreiner, Judy E. Bloom, Dylan S. Baker, Tiwanna M. Robinson, Richard Lieberman, Leslie M. Loew, Stormy J. Chamberlain, Eric S. Levine

**Author notes:** Corresponding Author: Eric S. Levine, Dept. of Neuroscience MC-3401, University of Connecticut School of Medicine 263 Farmington Ave., Farmington, CT 06030 USA.

## Abstract

Chromosome 15q11-q13 duplication syndrome (Dup15q) is a neurogenetic disorder caused by duplications of the maternal copy of this region. In addition to hypotonia, motor deficits, and language impairments, Dup15q patients commonly meet the criteria for autism spectrum disorder (ASD) and have a high prevalence of seizures. Here, we explored mechanisms of hyperexcitability in neurons derived from induced pluripotent stem cell (iPSC) lines from Dup15q patients. Maturation of resting membrane potential in Dup15q-derived neurons was similar to neurons from unaffected control subjects, but Dup15q neurons had delayed action potential maturation and increased synaptic event frequency and amplitude. Dup15q neurons also showed impairments in activity-dependent synaptic plasticity and homeostatic synaptic scaling. Finally, Dup15q neurons showed an increased frequency of spontaneous action potential firing compared to control neurons, in part due to disruption of KCNQ2 channels. Together these data point to multiple mechanisms underlying hyperexcitability that may provide new targets for the treatment of seizures and other phenotypes associated with Dup15q.

## Introduction

Chromosome 15q11-q13 is a region of the genome regulated by genomic imprinting. Most of the genes in this imprinted region are expressed exclusively from the paternally-inherited allele, while only one gene, *UBE3A,* is expressed exclusively from the maternally-inherited allele in mature neurons^1, 2^. Disruption of this genomic region can lead to three clinically distinct neurological disorders: Prader-Willi syndrome (PWS), Angelman syndrome (AS), and chromosome 15q11-q13 duplication syndrome (Dup15q)^3^. PWS is caused by the loss of the paternally-expressed genes on chromosome 15q11-q13^4^. Although AS is most commonly associated with large maternal deletions of 15q11-q13 encompassing many genes, it is known that loss of function from the maternal allele of *UBE3A* alone results in the full syndrome^5^. Dup15q results from duplications of maternal chromosome 15q11-q13^6^. Though the causative gene(s) for Dup15q is less clear, *UBE3A* is thought to play an important role in Dup15q pathophysiology. This is due to phenotypic similarities between AS and Dup15q, as well as the observation that individuals with maternal, but not paternal, duplications of chromosome 15q11-q13 have Dup15q and *UBE3A* is the only imprinted gene expressed from the maternally-inherited allele.

In addition to developmental delay, language impairments, and motor impairments, which are also commonly found in AS, Dup15q patients often meet the criteria for autism spectrum disorder (ASD)^7, 8^. In fact, Dup15q is the most common chromosomal anomaly associated with ASD^9^. Dup15q patients also have a high prevalence of seizures, which often do not respond well to commonly used medications^10^, and have an increased risk for sudden unexplained death in epilepsy (SUDEP)^11^. Investigations into cellular phenotypes of Dup15q neurons and their associated mechanistic underpinnings are important for the development of targeted therapeutics for seizures and other symptoms in these patients. Furthermore, understanding these cellular phenotypes may provide insight to the mechanisms causing idiopathic ASD.

To date, there are a variety of mouse models of Dup15q, but it is unclear how well they translate to the human Dup15q syndrome. For instance, one elegant mouse model was developed using chromosomal engineering to duplicate the entire chromosome 15q11-q13 syntenic region. While this mouse model has the best construct validity, disease relevant phenotypes are only observed when duplications are of paternal origin, rather than maternal origin^12^. BAC transgenics have also been used to express 2x and 3x copies of C-terminal tagged *UBE3A* to model Dup15q. Although these mice demonstrate behavioral phenotypes reminiscent of ASD, this model does not address the other genes that are duplicated in patients^13, 14^. Despite these shortcomings, it is very clear from the research with these mouse models that synaptic impairments are a strong component of Dup15q pathophysiology^12-16^. However, cellular phenotypes that may relate to seizures in these models are much less clear. The discovery of induced pluripotent stem cells (iPSCs) and the ability to differentiate these cells into neurons provides a unique opportunity to study human neurons with the exact genetic disruptions that cause Dup15q^17, 18^. Moreover, these neurons provide the ability to study the earliest stages of neuronal development, which is vital in the context of neurodevelopmental disorders such as Dup15q and ASDs.

We have previously identified a cellular phenotype in AS patient iPSC-derived neurons whereby synaptic and electrophysiological development, especially resting membrane potential (RMP), are impaired in these cells. This phenotype could also be recapitulated by either knocking down or knocking out *UBE3A* with antisense oligonucleotides or CRISPR/Cas9, respectively^19^. In the present study, we find that Dup15q neurons show a maturation of RMP that is similar to control neurons, though action potential (AP) development is slightly delayed, suggesting that the AS phenotype is specific to loss of *UBE3A* and not just indicative of neurodevelopmental disorders in general. Instead, Dup15q neurons show impairments in activity-dependent synaptic plasticity and homeostatic synaptic scaling. Additionally, Dup15q neurons have an elevated frequency of spontaneous AP firing as measured by both electrophysiology and calcium imaging. Finally, this AP firing phenotype seems to be due, at least in part, to disruption of the KCNQ2 potassium channel in these neurons, a channel linked to a genetic form of epilepsy. Together these data point to multiple mechanisms underlying hyperexcitability in Dup15q neurons, which may explain why seizures in Dup15q patients are difficult to treat. Moreover, the identification of these mechanisms of hyperexcitability may provide new targets for treatments and interventions for seizures and other phenotypes associated with Dup15q.

## Materials and methods

### Cell lines/Cell culture

iPSC lines from four Dup15q patients (1 male, 3 females), six unrelated unaffected controls (five males, one female), three AS patients (two males, one female), and one 15q11-q13 paternal duplication subject (female), were generated using either retroviral or lentiviral vectors expressing *OCT4, SOX2, KLF4, MYC, and LIN28*^45, 46^. iPSC lines from 3 Dup15q subjects (one male and two females) harboured an isodicentric chromosome of 15q11-q13. The fourth Dup15q patient (female) harbored an interstitial triplication of chromosome 15q11-q13. iPSC lines from two AS patients (one male and one female) harbored a large deletion of 15q11-q13. The third AS patient has a 2bp mutation to *UBE3A* (male patient)^47^. These iPSC lines and the female unaffected control were previously described and characterized^17^. Two of the Dup15q lines and the Pat Dup line were previously characterized^48^. Five control lines were generated from male donor subjects and have also been previously characterized^49, 50^. For all cell lines, patient samples were obtained under appropriate IRB protocols with consent. Cell culture experiments including stem cell plating, maintenance, and passaging, as well as neuronal differentiation and neuron maintenance and feeding were carried out as previously described^19^. Cell death assay was performed as previously described^19^.

### Electrophysiology

Whole-cell voltage and current clamp recordings were obtained from iPSC-derived neurons from Dup15q, AS, and 15q11-q13 paternal duplication patients and control subjects starting at 3 weeks post-initiation of differentiation (3 days post-plating). Electrophysiological measurements of intrinsic properties (Fig. 1) and synaptic development and plasticity (Fig. 2) were done as previously described^19^. For homeostatic synaptic plasticity experiments, culture media was treated with 1 µM TTX (Alomone Labs), 10 µM GABAzine (Sigma), or vehicle (aCSF) for 48 hours. Coverslips were then removed and placed in a recording chamber and continuously perfused with aCSF containing 1 µM TTX. Cells were held at -70 mV in voltage clamp for 1-3 minutes to monitor spontaneous excitatory synaptic currents. 2 coverslips per line per treatment were used. In all spontaneous firing experiments, cells were recorded at 25+ weeks in culture, 15 cells per coverslip and 1-2 coverslips per line per treatment were used. All cells were patched and RMP, evoked AP firing, and inward and outward currents were monitored. Following these protocols, all cells were then held at ∼-55 mV in current-clamp mode. Offline analysis was performed using Clampfit (Axon Instruments). All calcium imaging experiments were performed as previously described^19^. Tests of statistical significance were conducted using analysis of variance (ANOVA), Student’s t-test, Kolmogorov-Smirnov, or the chi-square test as noted. Data are presented as mean ± standard error of the mean.

**Figure 1.**
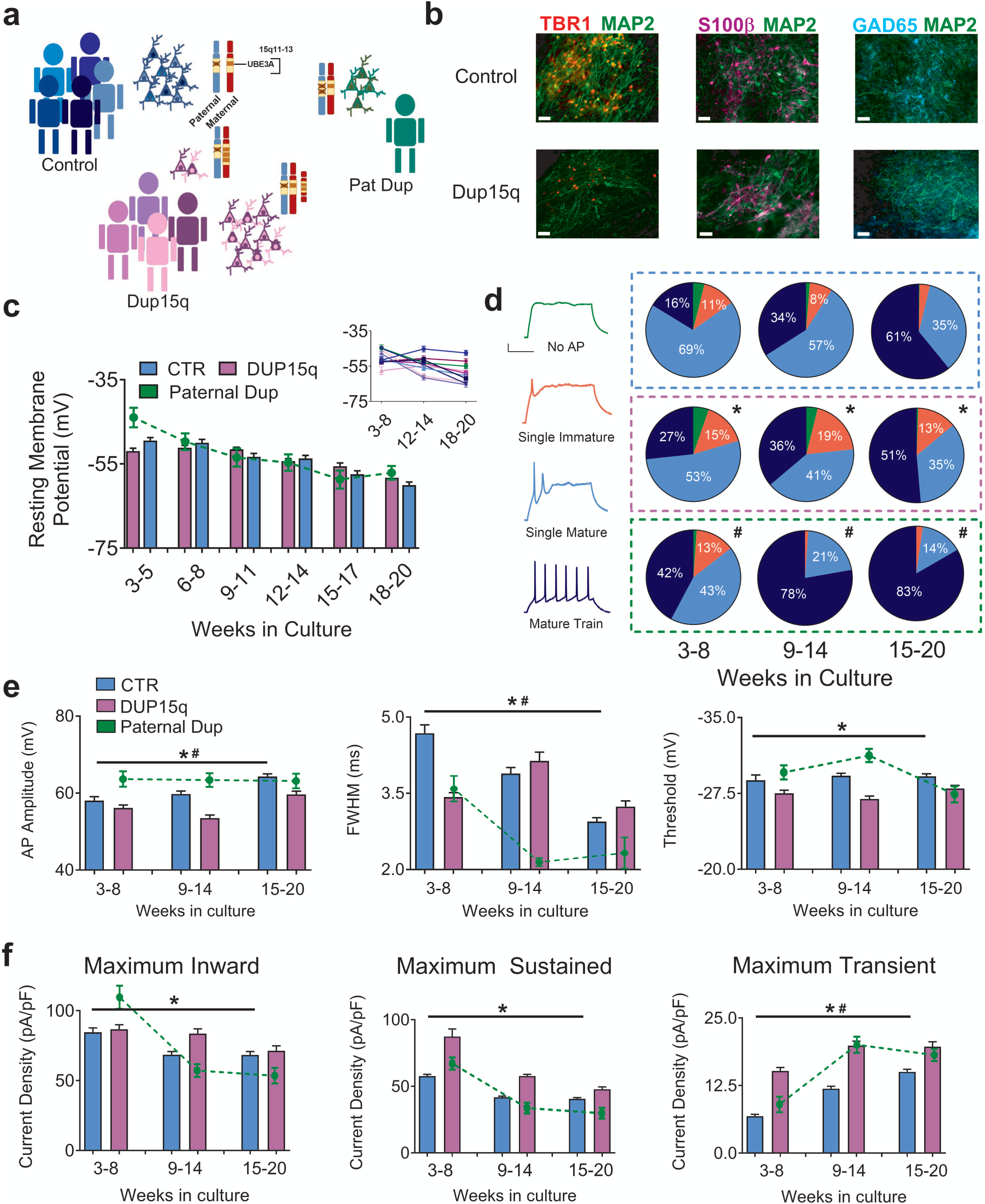
Electrophysiological profile of iPSC-derived neurons from control and Dup15q subjects during *in vitro* development. (**a**) Schematic of patient line genetics of the chromosome 15q11-13 genomic locus. (**b**) Immunocytochemical staining for TBR1/MAP2, S100β/MAP2, and GAD65/MAP2, in control and Dup15q-derived cultures. Scale bar, 50 µm (**c**) Group data for resting membrane potential (RMP) of control (CTR; 5 subjects; n>200 at each time point), Dup15q (4 subjects; n>175 at each time point) and a subject with a paternal duplication of chromosome 15q11-q13 (Paternal Dup; 1 subject; n=45 at each time point) during development. Inset: RMP for all individual lines. For each time bin, n≥30 for all lines. (**d**) Left: Example traces representing four AP firing patterns used for characterization. Scale bar, 20 mV, 200 ms. Right: Distribution of AP firing patterns for control (blue box; 5 subjects; n>430 at each time point), Dup15q (purple box; 4 subjects; n>430 at each time point), and a 15q11-q13 paternal duplication (green box; 1 subject; n>60 at each time point) neurons at three developmental time bins. *P<0.0001 for differences between control and Dup15q; χ^2^ test. #P<0.0001 for differences between control and paternal duplication; χ^2^ test. (**e**) AP amplitude (left), full width at half-maximum amplitude (FWHM; middle), and AP threshold (right), for control, Dup15q, and paternal duplication cultures at three time points (n>350 for both control and dup15q at all time points; n>80 for paternal duplication at all time points). *P<0.001 for significant differences between control and Dup15q, ^#^P<0.001 for significant differences between control and paternal duplication (two-way ANOVA). (**f**) Group data for maximum inward current density (left), maximum sustained outward current density (middle), and maximum transient outward current density (right), for control, Dup15q, and paternal duplication cultures at three time points (n>350 for both control and dup15q at all time points; n>80 for paternal duplication at all time points). *P<0.001 for significant differences between control and Dup15q, ^#^P<0.001 for significant differences between control and paternal duplication (two-way ANOVA).

**Figure 2.**
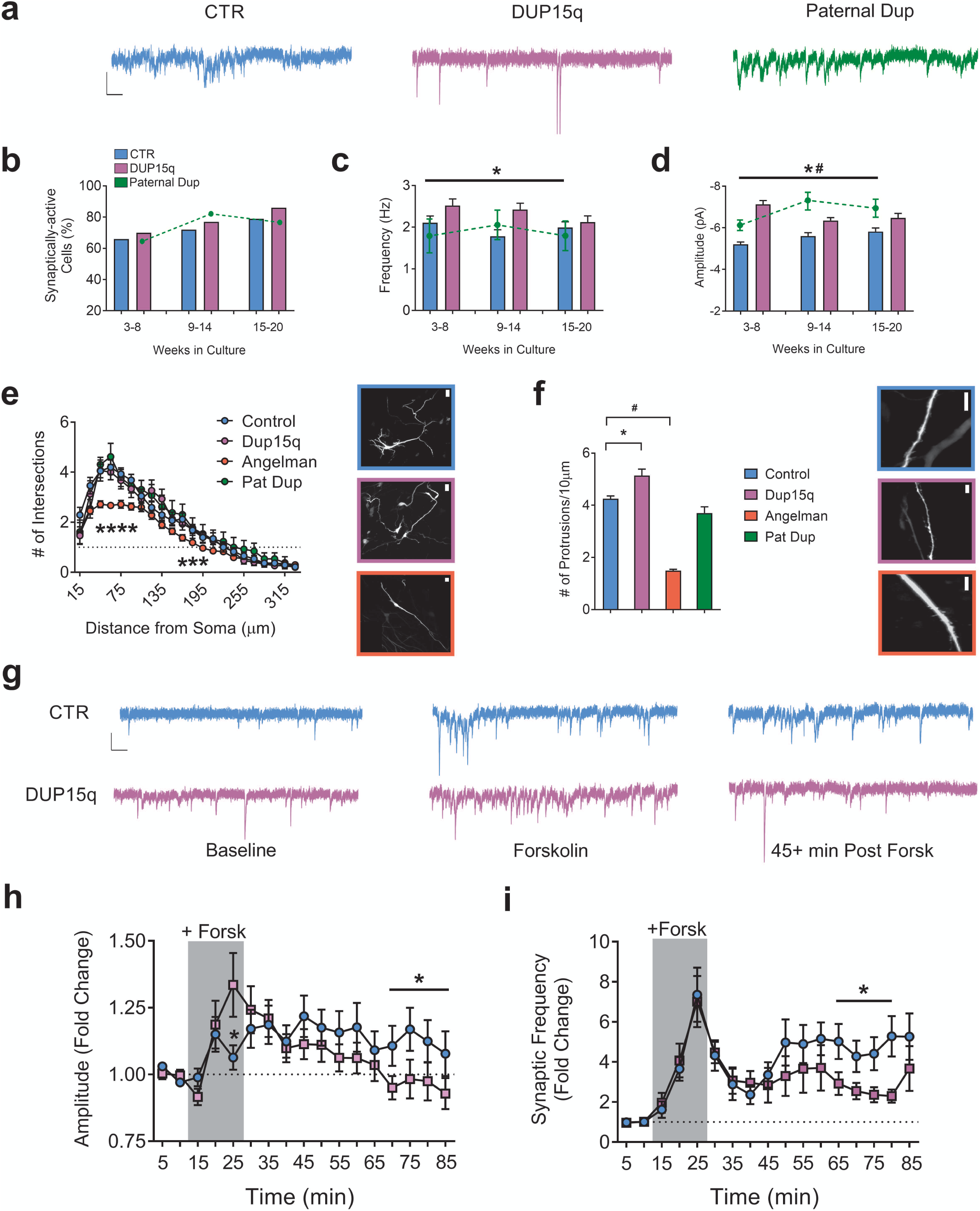
Development of synaptic activity and expression of activity-dependent synaptic plasticity. (**a**) Example traces of spontaneous excitatory synaptic currents from control (CTR), Dup15q, and 15q11-13 paternal duplication (Paternal Dup) neurons at 15-20 weeks in culture. Scale bar: 10 pA, 100 ms. (**b**) Percent of synaptically-active neurons (event frequency at >0.2 Hz) derived from CTR (5 subjects; n>415 at every time point), Dup15q (4 subjects; n>350 at every time point), and paternal duplication (1 subject; n>80 at every time point). *P<0.05 for significant differences between control and Dup15q. ^#^P<0.05 for significant differences between control and paternal duplication (χ^2^ test). (**c**) Mean frequency of spontaneous synaptic events for active neurons derived from CTR (5 subjects, n>250 cells at every time point), Dup15q (4 subjects, n>315 cells at every time point) and 15q11-13 paternal duplication (1 line; n>50 cells at every time point). (**d**) Mean amplitude of spontaneous synaptic events for neurons plotted in (**c**). *P<0.05 for differences between CTR and AS. ^#^P<0.05 for differences between CTR and paternal dup (two-way ANOVA). (**e**) Left: Grouped Sholl analysis for neurons (28-30 weeks) from control-(n>20), Dup15q-(n=11), AS-(n>40), and 15q11-q13 paternal duplication-derived (n=5) cultures. *P<0.05 indicated significant differences between control and AS (Student’s t-test). Right: example images of transfected neurons used for analysis from control (top; blue), Dup15q (purple; middle), and AS (red; bottom). Scale bar: 25µm. (**f**) Left: Grouped analysis for number of dendritic protrusions for control (n>45), Dup15q (n=20), AS (n>40), and 15q11-13 paternal duplication cultures (n=9). Data from weeks 15-20 and 28-30 were collapsed into a single bin. *P<0.05 indicates significant differences between control and Dup15q. ^#^P<0.05 indicates significant differences between control and AS (Student’s t-test). Right: example images of transfected neurons used for analysis from control (top; blue), Dup15q (purple; middle), and AS (red; bottom). Scale bar: 10µm. (**g**) Example traces of spontaneous excitatory synaptic currents from control (CTR; top) and Dup15q (bottom) neurons at (left to right) baseline, during forskolin/rolipram/0 Mg (Forsk), and 45+ min post-Forsk (see Methods for details). Scale bar, 10 pA, 100 ms. (**h,i**) Group data for CTR (n>60) and Dup15q (n>45) neurons showing (**h**) mean frequency and (**i**) mean amplitude of spontaneous synaptic currents during baseline, plasticity induction (indicated by grey box; see Methods for details) and post induction. *P<0.05, indicates differences for CTR vs Dup15q (Student’s t-test).

### Live-cell image acquisition

iPSC-derived neuronal cultures were transfected using Lipofectamine 3000 (ThermoFisher) and the plasmid FUGW 51. FUGW was a gift from David Baltimore (Addgene plasmid #14883). Neurons were imaged 48 hours after transfection. Coverslips were inverted and placed in imaging chamber RX-26G (Harvard Apparatus) with 500 µl of Live Cell Imaging Solution (ThermoFisher). Neuron were imaged on Zeiss 780 confocal microscope with, 63x oil-immersion objective with a NA of 1.4 and 3x zoom. Neurons with clearly distinguishable processes were imaged as a Z-series (0.2μm interval) and tile scan to capture the entire cell. At least 3 randomly chosen neurons were imaged for each cell line, a minimum of 9 neurons were used for each phenotype for both time points (18-20 weeks: Control 27 cells, Dup15q 9 cells; 28-30 weeks: Control: 21 cells, Dup15q 9 cells, AS 42 cells). Maximum projection-stitched images were created using Zeiss ZEN Microscope Software.

### Quantitative image analysis

NIH ImageJ software^53^ was used for both Sholl analysis and protrusion quantification. Image analysis was done blinded to experimental condition. Semiautomatic tracings of individual cells were acquired using the Simple Neurite Tracer plugin^54^ for ImageJ, and tracing files were generated. The trace files were then used for Sholl analysis using the Sholl Analysis plugin^55^. Protrusion quantification was done manually, protrusions > 2 µm are likely to be filopodia and were not included in the analysis. Quantification began 50 µm from the soma to avoid large variations in protrusion densities between cells^56^ and continued the entire length of the dendrite. In order to detect all protrusions that may be hidden in a collapsed z-stack, individual planes were examined. Density was calculated by quantifying the total number of protrusions per entire dendritic length and expressed as protrusions/10 μm.

### Immunocytochemistry

For immunocytochemical experiments, iPSC-derived neurons were fixed in 4% paraformaldehyde in PBS for 15 min. Cell cultures were washed and permeabilized with TNT buffer for 20 min at 4**°** C. Cultures were then blocked with TNB buffer for 30 minutes at 4 °C and incubated overnight at 4° C in primary antibodies (in combinations as indicated) against previously described antibodies^19^. the following morning and counter stained with secondary antibody diluted in TNB buffer (Perkin Elmer, TSA blocking reagent, FP1020) for 2 hours at room temperature. Secondary antibodies used were previously described^19^. Following secondary staining, cells were washed with TNT buffer and stained with DAPI (Sigma 10236276001) for 7 minutes. After staining procedures were complete coverslips were mounted onto slides with PPD (Sigma P23962/P6001) + Mowial 4-88 (Sigma 81381-50G). Slides were imaged with a Zeiss Axiovert.

### Flow cytometry

Flow cytometry was performed as previously described^19^. Primary antibodies and concentrations used were MAP2 (mouse anti-human, 1:500, Sigma M 1406), KCNQ2 (rabbit anti-human, 1:200, Sigma AV35459) and chicken anti-human TUJ1 (Covance or Abcam, 1:500). The following day cells were washed with FACS buffer and incubated with secondary antibody (anti-mouse Alexafluor 488 1:1000, anti-rabbit Alexafluor 594 1:1000, goat anti-chicken Alexafluor 647 1:1000) at 4**°** C for one hour. Cells were then analyzed on a BD Biosciences LSRII flow cytometer.

## Results

### Electrophysiological characterization of Dup15q and control cultures

iPSC lines were generated and differentiated into neurons as previously described from five unaffected control (CTR) subjects, four chromosome 15q duplication (Dup15q) patients, and a single individual with a paternal chromosome 15q11-q13 duplication who did not have Dup15q or ASD (Pat Dup) (Fig. S1a; Fig. 1a)^20^. Given that *UBE3A* is paternally silenced in neurons, the Pat Dup line provides a tool for examining the involvement of overexpression of genes other than *UBE3A* housed within 15q11-13, including GABAR subunits. Neurons were also generated from three Angelman syndrome (AS) patients (Fig. S1b). The majority of cells produced by our differentiation protocol express the neuronal marker MAP2 and the glutamatergic marker TBR1 (70-80%)^19^, with a small percentage of cells expressing the GABAergic marker GAD65 or the astrocyte marker S100β. The proportion of these cell types is not qualitatively different between CTR and Dup15q-derived cultures (Fig. 1b) and is similar to our previously published data in control and AS-derived cultures^19^.

We recently discovered a robust electrophysiological phenotype in iPSC-derived neurons from AS patients whereby these neurons show an impairment in the maturation of RMP, AP firing, and synaptic activity. Moreover, these phenotypes were the specific result of loss of *UBE3A*, the gene responsible for AS, as knockdown or knockout of *UBE3A* could recapitulate the entire spectrum of developmental deficits^19^. Given the close association of AS and Dup15q and the shared involvement of *UBE3A* in these syndromes, we examined electrophysiological development of patient-derived Dup15q neurons. Whole-cell patch-clamp recordings were performed on ∼15 cells/coverslip/week/line starting at plating (week 3) through 20 weeks in culture. Current clamp and voltage clamp protocols were used on all cells to measure RMP, AP firing, inward and outward voltage-gated currents, and synaptic activity. As expected, neurons derived from control subjects show a shift to more hyperpolarized RMP values across 20 weeks in culture (Fig. 1c). However, unlike AS-derived neurons, Dup15q and paternal dup neurons show a normal development of RMP, suggesting that this impairment is specific to loss of *UBE3A*. Results from individual lines are also shown (Fig. 1c).

We next categorized AP firing in neurons during *in vitro* development. Neurons were held at ∼-70 mV in current-clamp mode and 5 pA current steps from -20 to +40 were applied to each cell. APs were classified as either No AP, Single Immature, Single Mature, or Mature Train (Fig. 1d; left). In control neurons, a variety of AP firing types are initially seen, switching to predominantly mature firing as development proceeds (Fig. 1d; right). Interestingly, although they do not show the developmental disruption in RMP observed in AS neurons, Dup15q-derived neurons show a delay in AP maturation compared to controls. Conversely, recordings from the Pat Dup line show an enhanced maturation of AP firing compared to controls, however only a single Pat Dup cell line was investigated.

AP development was further characterized by examining amplitude, full-width at half maximum (FWHM), and threshold, as well as inward and outward currents that correspond to different phases of the AP. In support of the AP firing data, control neurons show an increase in AP amplitude, a decrease in FWHM, and an increase in transient outward current during *in vitro* development (Fig. 1e,f), indicative of AP maturation. These results are consistent with our previously published data on 4 separate control lines^19^. Dup15q-derived neurons have a significantly decreased AP amplitude and increased FWHM across development compared to controls, which is also representative of the more immature AP firing observed in Fig. 1d. Pat Dup neurons have a significantly increased AP amplitude and decreased FWHM across development compared to controls. Additionally, Dup15q neurons show significant increases in maximum inward, sustained outward, and transient outward currents compared to control neurons, which may also account for the differences observed in AP firing (Fig. 1e,f). Cell capacitance, which increased during *in vitro* development, was not different between CTR and Dup15q neurons (Fig. S1c). Importantly, cell death, as measured by an LDH assay, was also not significantly different between control and Dup15q cultures (Fig. S1d).

### Spontaneous synaptic transmission and synaptic plasticity

Both deletion and duplication of *UBE3A* have been closely linked to synaptic impairments^13-15, 21, 22^. Furthermore, loss of *UBE3A* in mouse models of AS leads to disrupted long-term potentiation and depression^19, 23-26^. We have observed similar phenotypes in iPSC-derived neurons from AS patients^19^. For this reason, we measured the development of spontaneous excitatory synaptic activity in control, Dup15q, and Pat Dup cultures. Example traces of spontaneous activity are depicted in Fig. 2a. Briefly, neurons were held at -70 mV for 1 to 3 minutes and synaptic events were detected and quantified as described (See Methods). Overall, control, Dup15q and Pat Dup cultures showed similar percentages of synaptically-active cells, which gradually increased in all three genotypes across development (Fig. 2b). Both the frequency and amplitude of spontaneous synaptic events from synaptically-active cells were significantly increased in Dup15q neurons compared to controls (Fig. 2c,d). Interestingly, amplitude of synaptic events was also significantly increased in Pat Dup cells (Fig. 2d).

We next examined whether morphological changes were associated with the synaptic phenotype in Dup15q neurons. Briefly, neurons were transfected via Lipofectamine 3000 with a plasmid with hUbC-driven EGFP and imaged 48 hours after transfection. Images were collected at either 15-20 weeks or 28-30 weeks and Sholl analysis was performed to quantify dendritic branching (See Methods). Neither Dup15q nor Pat Dup neurons showed any differences in dendritic complexity at either 15-20 weeks (Fig. S2a) or 28-30 weeks (Fig. 2e) as compared to control. As we have previously shown that AS-derived neurons have decreased synaptic activity^19^, we also performed Sholl analysis on 3 AS-derived patient lines. Unlike Dup15q and Pat Dup neurons, AS neurons had a significant decrease in dendritic complexity compared to controls (Fig. 2e). Example images of transfected neurons from control, Dup15q, and AS cultures are shown in Fig. 2e (right). Additional examples and neuron traces are depicted in Fig. S2b,c. Sholl analysis for 15-20 week old cultures and for individual lines are also depicted in Fig. S2a,c. We next counted the number of protrusions along the dendritic tree. Dup15q neurons showed a significant increase in the number of protrusions and AS neurons showed a significant decrease in the number of protrusions, compared to controls (Fig. 2f). Pat Dup neurons were not significantly different from controls. Thus, duplications and deletions of 15q11-q13 result in increases and decreases, respectively, in the number of dendritic protrusions and frequency of excitatory synaptic events in iPSC-derived cultures. Example images of dendritic protrusions from transfected control, Dup15q, and AS neurons are depicted in Fig. 2f (right). Number of protrusions for individual control, Dup15q, and AS lines are shown in Fig. S2d. We have previously shown that iPSC-derived neurons can undergo activity-dependent synaptic plasticity as measured by prolonged increase in spontaneous synaptic event frequency following exposure to a cocktail of forskolin and rolipram in a 0 Mg^2+^ solution, which simultaneously boosts cAMP levels and enhances NMDA receptor activation^19, 27^. Using this protocol, AS-derived neurons showed impairments in long-term plasticity compared to controls. We used this same protocol to measure activity-dependent synaptic plasticity in Dup15q and additional control cultures. Both control and Dup15q neurons showed an increase in synaptic event amplitude and frequency over the course of the fifteen-minute induction period (Fig. 2h,i; grey box; See Methods). Following washout, control neurons maintained a significant increase in both spontaneous synaptic amplitude (Fig. 2h) and frequency (Fig. 2i) across the entire one-hour of post-induction recording. Though Dup15q neurons showed similar maintained increases in amplitude and frequency early post-induction (five to forty minutes), they showed significantly diminished plasticity of both measures for the remainder of the post-induction period compared to controls.

### Disrupted synaptic scaling in Dup15q neurons

In mouse models, deletion or overexpression of *UBE3A* causes changes in the expression of the protein Arc, though these changes do not seem to be related to UBE3A’s role as an ubiquitin ligase^13, 28-31^. Arc is a major synaptic protein involved in the trafficking of AMPA receptors at the synapse and modulating synaptic strength^32, 33^. Given this correlation and the synaptic deficits related to various models of AS and Dup15q, we next looked at homeostatic synaptic plasticity/synaptic scaling, a well-established phenomenon in which temporarily changing baseline levels of network activity in neurons results in opposite changes in synaptic activity that are the result of increases or decreases in the number of AMPA receptors at the synapse^34, 35^. Importantly, this type of plasticity has been shown to exist in cultured iPSC-derived neurons and is disrupted in a mouse model of AS^29, 36^. We therefore measured the amplitude of AMPA receptor-mediated miniature excitatory postsynaptic currents (mEPSCs) subsequent to GABAzine or TTX treatment. Following disinhibition with GABAzine, control neurons showed a leftward shift in the amplitude distribution that was not observed in Dup15q neurons (Fig. 3a). Due to line-to-line variability, data from each line was normalized to its own baseline distribution and then grouped by genotype. Example traces are depicted in Fig.3b. Interestingly, unlike Dup15q neurons, but similar to controls, AS neurons showed a decrease in AMPA event amplitude following GABAzine treatment. To our surprise, all genotypes failed to respond to TTX with a rightward shift in AMPA event amplitude. We next asked whether a low baseline frequency of spontaneous mEPSCs might occlude the effect of TTX. In support of this, we found that 2 control lines with mean baseline activity >3 Hz did in fact display the expected rightward shift in AMPA current amplitudes with TTX treatment (Fig. 3c). None of the Dup15q cultures met the 3 Hz criteria. Additionally, we compared the baseline (vehicle-treated) amplitude distributions of control, Dup15q, and AS neurons. In line with our results in Fig. 2d for Dup15q neurons and our previously published data on AS-derived neurons, both AS and Dup15q baseline amplitudes were significantly increased compared to control (Fig. 3c). These data may be explained in part by the inability of Dup15q neurons to decrease mEPSC amplitudes in response to increases in network activity.

**Figure 3.**
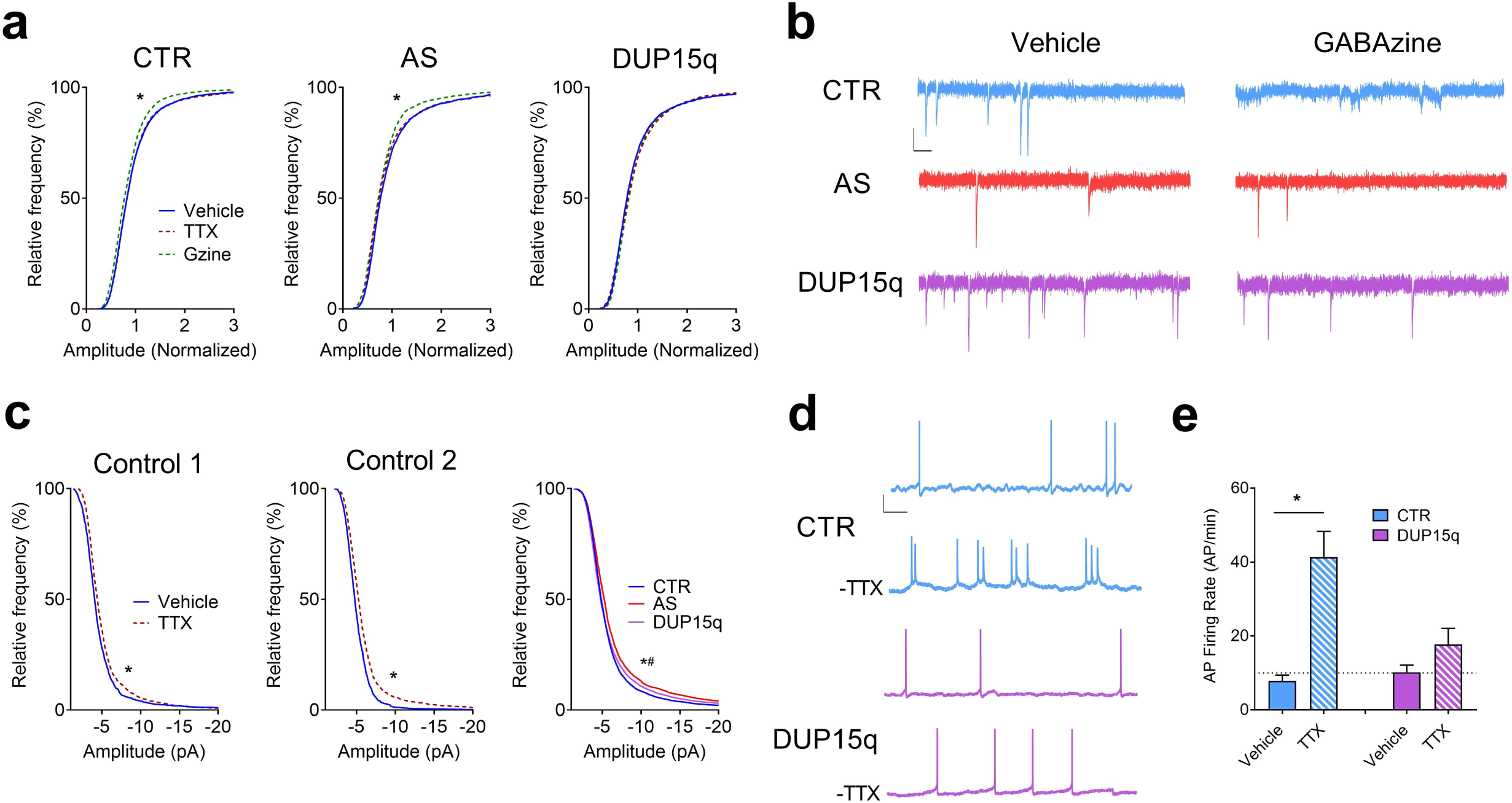
Impaired synaptic scaling in Dup15q-derived neurons. (**a**) Cumulative frequency histograms of amplitudes (normalized) of miniature spontaneous excitatory synaptic currents for control (CTR; left; 6 subjects; n>140 neurons per treatment), Angelman (AS; middle; 4 subjects; n>80 neurons per treatment), and Dup15q (right; 4 subjects; n>100 neurons per treatment) cultures. *P<0.0001 indicated significant differences between control and Gzine treatments (Student’s t-test). (**b**) Example traces of miniature spontaneous excitatory synaptic currents from CTR, AS, and Dup15q neurons following treatment with either vehicle (left) or GABAzine (right). Scale bar: 10 pA, 100ms. (**c**) Cumulative frequency histograms of raw amplitudes of miniature spontaneous excitatory synaptic currents for 2 control subjects treated with TTX (Control 1; left. Control 2; middle), with baseline synaptic frequencies >3 Hz. Right: Cumulative frequency histograms of raw baseline (vehicle-treated) amplitudes for control, AS, and Dup15q cultures. *P<0.0001 indicates significant differences between control and Dup15q, #P<0.0001 indicates significant differences between control and AS (Kolmogorov-Smirnov test). (**d**) Example traces of spontaneous action potential firing from control (CTR) and Dup15q neurons from data presented in (**e**). Scale bar: 20 mV, 1s. (**e**) Spontaneous action potential (AP) firing rate of vehicle- or TTX-treated neuronal cultures from control and Dup15q subjects following washout of TTX. *P<0.05 indicates significant differences between vehicle- and TTX-treated cultures (Student’s t-test).

It is thought that homeostatic synaptic scaling allows a neuron to maintain a set-point of AP firing in response to changes in network synaptic activity and synaptic strength^37, 38^. Thus, the frequency of AP firing should change in the same direction as AMPA currents following treatment with TTX. For this reason, we recorded spontaneous AP firing (following washout) in control and Dup15q neurons that had been treated with either vehicle or TTX for 48 hours. Example traces of spontaneous APs are depicted in Fig. 3d. As expected, control neurons responded to TTX treatment with a significant increase in spontaneous AP firing (Fig. 3e), whereas Dup15q neurons failed to show a significant increase in AP firing. Overall, these results suggest a disruption in the ability of Dup15q neurons to scale AMPA-mediated synaptic currents in response to changes in network activity. This deficit, particularly the inability to down-regulate synaptic currents in response to increased network activity, suggests a hyperexcitable phenotype that could contribute to the high prevalence of seizures in Dup15q patients.

### Spontaneous AP firing

We next recorded spontaneous AP firing from 9 control, 4 Dup15q, 1 Pat Dup, and 3 AS lines at 25+ weeks in culture. Neurons were held at rest (∼-55 mV) in current-clamp mode to measure spontaneous firing. Example traces are shown in Fig. 4a. The frequency of spontaneous APs recorded from individual lines was slightly variable, but was consistent within each genotype (Fig. 4b). On average, control neurons fired less than 0.5 Hz, which was similar to Pat Dup and AS-derived cultures. Dup15q neurons, however, had almost triple the firing rate of the other genotypes (Fig. 4c). The amplitudes of spontaneous APs were not different between genotypes (Fig. 4d, e). We also indirectly monitored AP firing in multiple neurons simultaneously using calcium imaging. Calcium transients were quantified over the course of a 10-minute imaging period and plotted as Raster plots (Fig. 4g). Consistent with patch-clamp recordings, cultures from 2 Dup15q patients showed a higher frequency of calcium transients and, in both cases, Dup15q neurons showed a high degree of synchrony, which was not observed in the control cultures. This was also true for even the most active control neurons from an additional experiment (See example traces in Fig. 4f). Together these data provide an additional aspect of hyperexcitablity that may relate to the frequent seizures in Dup15q patients.

**Figure 4.**
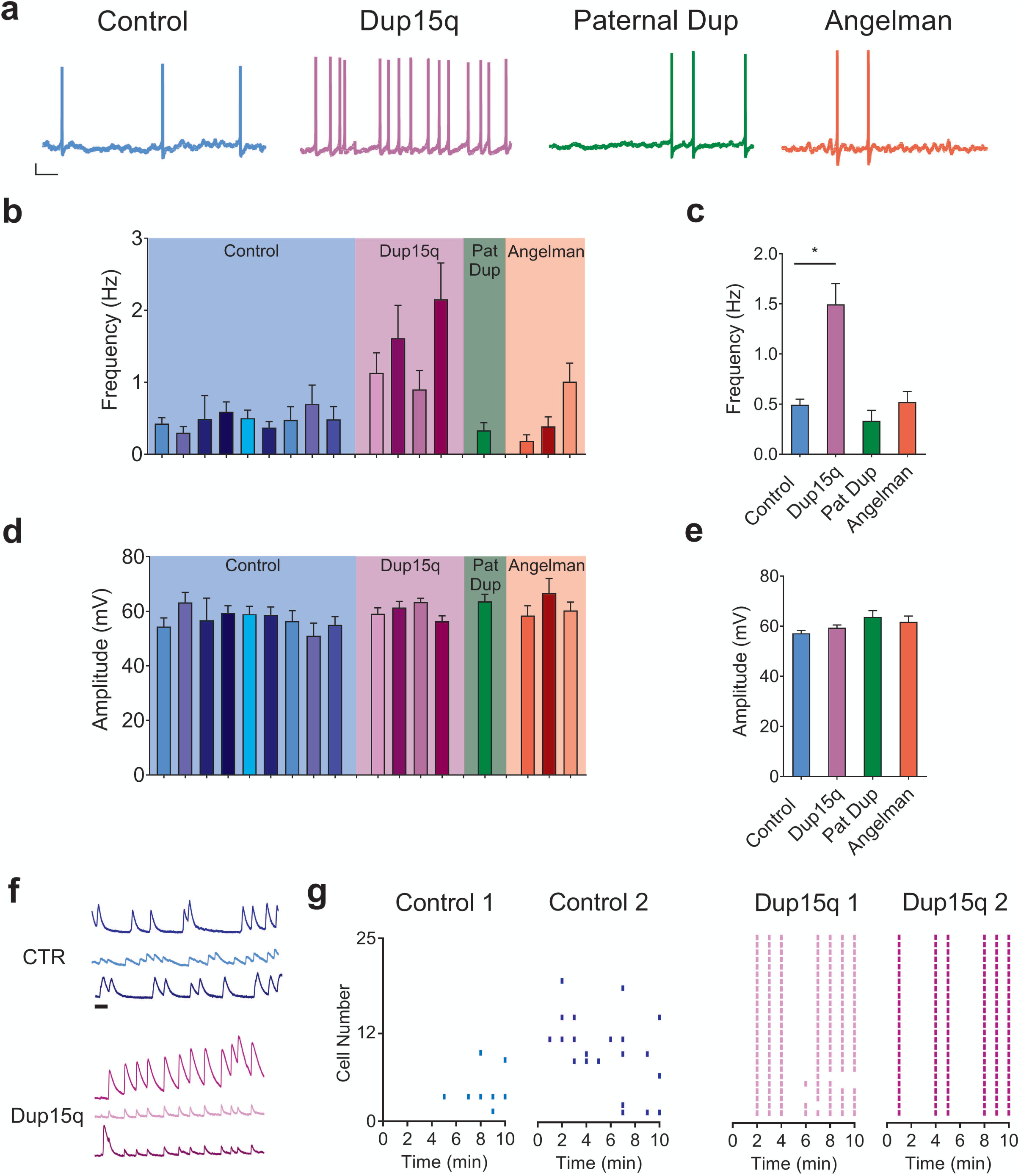
Spontaneous action potential firing. (**a**) Example traces of spontaneous action potential firing from control (CTR; blue), Dup15q (purple), 15q11-13 paternal duplication (Paternal Dup; green) and AS (red) neurons at 25+ weeks in culture. Scale bar: 10 mV, 1s. (**b,d**) Spontaneous action potential frequency (**b**) and amplitude (**d**) of neurons derived from individual control lines (blue; 9 subjects; n>15 neurons per bar), Dup15q lines (purple; 4 subjects; n>15 neurons per bar), 15q11-q13 paternal duplication (green; 1 subject; n>25), and AS (red; 3 subjects; n>25 per bar). (**c,e**) Grouped data of data presented in (**b,d**) (n=224, n=110, n=32, n=89 for control, Dup15q, Pat Dup, and AS, respectively). *P<.05 indicates significant differences between control and Dup15q (Student’s t-test). (**f**) Example traces of spontaneous baseline calcium transients from 3 neurons each from CTR (top; blue) and Dup15q (bottom; purple). Scale bar, 1 min. (**g**) Raster plots of frequency of calcium transients from cultures derived from control (left; 2 subjects; 25 neurons each) and Dup15q (right; 2 subjects; 25 neurons each).

### KCNQ2 channels are disrupted in Dup15q neurons

Given the role of KCNQ2 potassium channels as a brake on repetitive neuronal AP firing and its involvement in epilepsy early in development, we decided to further explore potential KCNQ2 deficits in Dup15q-derived neurons. Recordings were made from 15 cells from 1-2 coverslips from 9 control subjects, 4 Dup15q subjects, 3 AS subjects, and 1 Pat Dup subject in the presence of either normal ACSF, KCNQ2 blocker XE991, or KCNQ activator flupirtine. As with experiments in Fig. 4, neurons were held in current-clamp at rest (∼-55mV). Example data of spontaneous AP frequency from 3 individual control (Fig. 5a) and 3 individual Dup15q (Fig. 5b) lines are shown. Additional data from individual lines are also shown in Fig. S3a. Example traces from control and Dup15q neurons during the 3 conditions are depicted in Fig. 5c,d. Under baseline conditions, control neurons fire spontaneous APs at a frequency of ∼0.5 Hz (Fig. 5e). As expected, control neurons recorded in the presence of the KCNQ2 activator flupirtine have a significantly reduced frequency of AP firing. In the presence of the KCNQ2 blocker XE991, control neurons have a significantly increased level of spontaneous AP firing that is strikingly similar to the baseline level of spontaneous AP firing in Dup15q neurons (Fig. 5e). Both Pat Dup and AS neurons in the presence of either flupirtine or XE991 responded similarly to controls. Dup15q neurons, however, failed to respond to either drug, suggesting an impairment in KCNQ2 function in these cells. AP amplitude was unaffected across genotype groups and pharmacologic treatment (Fig. 5f). Additional AP amplitude data from individual lines is shown in Fig. S3b. To better understand the lack of response to pharmacological manipulation of KCNQ2 in Dup15q neurons, we stained control and Dup15q cultures for MAP2 and KCNQ2 and then sorted cultures by flow cytometry. We found that >60% of control MAP2+ cells were also positive for KCNQ2 (Fig. 5g), and that this percentage was significantly less in Dup15q cultures (∼25%), providing evidence for decreased KCNQ2 expression in Dup15q neurons.

**Figure 5.**
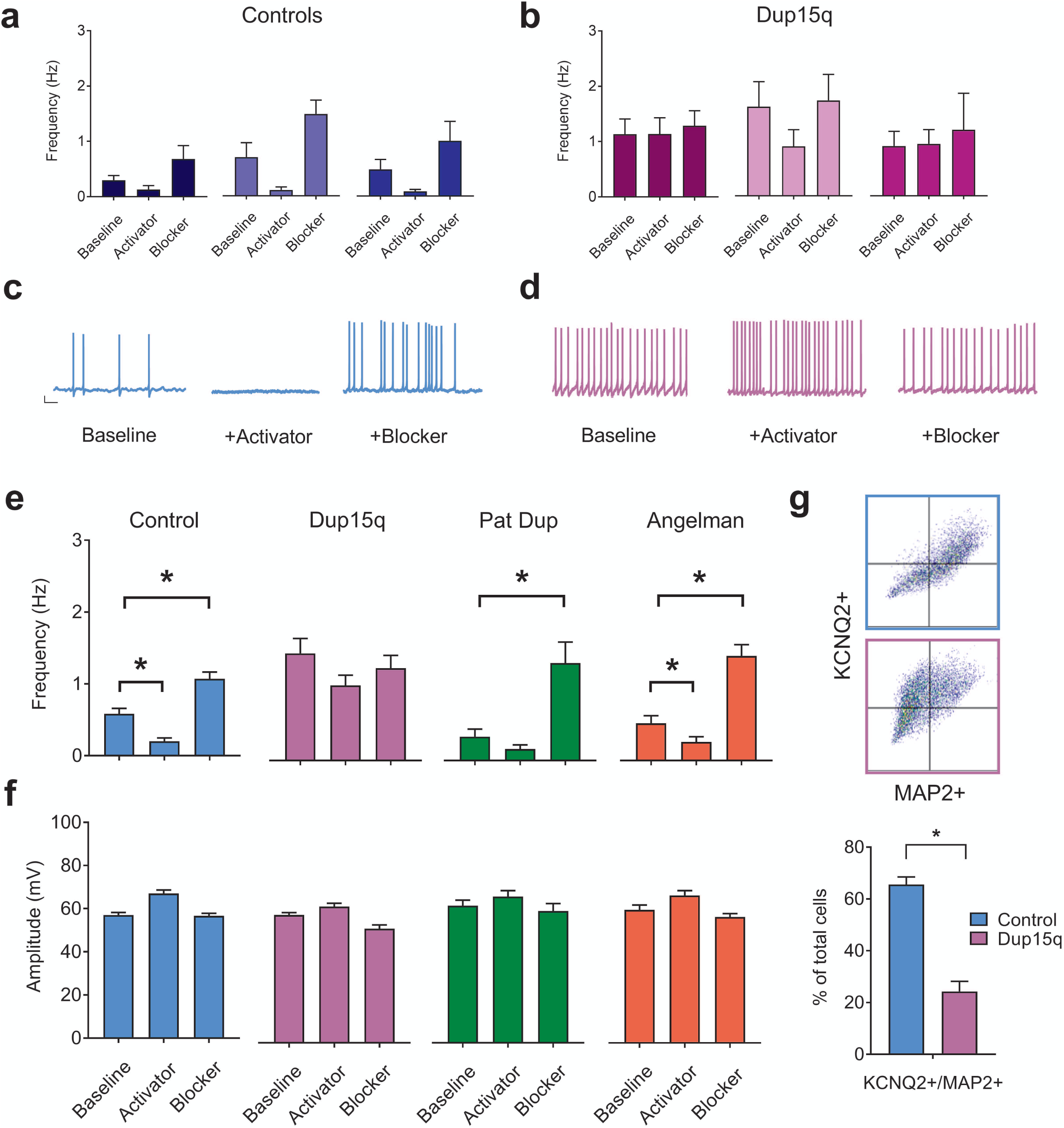
Impaired KCNQ2 channels in Dup15q-derived neurons. (**a,b**) Frequency of spontaneous action potential firing for 3 individual control (**a**) and Dup15q (**b**) lines at baseline or in the presence of either pharmacological blocker or activator of KCNQ2 channels (n>15 neurons for each bar).(**c**) Example traces of spontaneous action potentials recorded from control neurons. Scale bar: 10mV, 1s (**d**) Example traces of spontaneous action potentials recorded from Dup15q neurons (**e**) Group data for frequency of spontaneous action potential firing for control (left; blue; 9 lines; n=251, 160, 204 neurons for baseline, activator, and blocker, respectively), Dup15q (middle left; purple; 4 lines; n=110, 108, 93 neurons for baseline, activator, and blocker, respectively), 15q11-13 paternal duplication (middle right; green; 1 line; n=30 neurons for all bars), and Angelman (right; red; 3 lines; n>89 neurons for all bars), recorded at baseline or in the presence of either pharmacological blockers or activators of KCNQ2 channels. *P<0.05 indicates significant differences (student’s t-test). (**f**) Group data for amplitude of spontaneous action potential firing for data presented in (**e**). (**g**) Top: Examples of flow cytometry plots for cultures stained with KCNQ2 and MAP2. bottom: Flow cytometry quantification of percent of cells positive for both MAP2 and KCNQ2 for control (2 subjects/8 coverslips) and Dup15q (2 subjects/8 coverslips).*P<0.05 indicates significant differences (Student’s t-test).

## DISCUSSION

Deletions or duplications of maternal chromosome 15q11-q13 result in AS or Dup15q, respectively, distinct syndromes that share some clinical phenotypes including a high prevalence of seizures^7^. We have previously shown that iPSC-derived neurons from AS patients have a more depolarized RMP compared to controls across 20 weeks of *in vitro* development, which was the result of loss of *UBE3A*, a gene housed in the 15q11-q13 region that encodes for an ubiquitin ligase protein and is known to be the causative gene for AS^19^. A depolarized RMP in AS neurons represents a mechanism of hyperexcitability in these cells that could relate to the seizures in AS patients. In addition, human AS neurons showed deficits in AP development and synaptic transmission. In the present study, we aimed to test whether a similar cellular phenotype is found in Dup15q syndrome patient-derived neurons due to the presence of *UBE3A* in the duplicated region. Interestingly, RMP development in Dup15q neurons was similar to controls over 20 weeks of *in vitro* development, shifting to more hyperpolarized potentials over time in culture. Development of AP firing was delayed in Dup15q neurons compared to controls, but was not as strongly impaired as observed in AS neurons. Together, these data suggest separate and distinct mechanisms of hyperexcitability for Dup15q patient neurons.

Mouse models of both AS and Dup15q strongly indicate synaptic dysfunction as an important contributor to pathophysiology in these syndromes^13, 14, 22, 39^. Here, we observe similar deficits in synaptic function in Dup15q patient-derived neurons. First, Dup15q neurons show significant increases in synaptic event frequency and amplitude compared to controls, which is maintained over 20 weeks of *in vitro* development, but no differences in dendritic branching. Similar increases in synaptic event amplitude have been observed in AS-derived cultures, though these cells show significant decreases in synaptic frequency. Second, although control neurons display a long-term increase in the frequency and amplitude of synaptic events in response to pharmacologically-induced plasticity, Dup15q neurons fail to show similar long-term plasticity.

Increases and decreases in *UBE3A* have previously been linked to changes to the synaptic protein Arc, a molecule responsible for internalization of AMPA receptors at the synapse ^13, 29, 30, 40, 41^. In line with these data and the observed increase in amplitude of synaptic events across development, we find that Dup15q neurons fail to display homeostatic synaptic scaling in response to either disinhibition or Na^+^-channel block, which is normal in both control and AS cultures. These differences were reflected in both synaptic and spontaneous AP firing responses. The goal of homeostatic scaling and plasticity is to establish a mechanism to avoid positive-feedback loops that continually increase AP firing in response to long-term potentiation. Therefore, an impaired ability for Dup15q neurons to decrease synaptic amplitudes in response to changes in network activity represents a potential mechanism for hyperexcitability in these neurons.

Given the seizure phenotype commonly associated with Dup15q, we also measured spontaneous AP firing in control, AS, and Dup15q neurons. Control and AS neurons showed low levels of baseline spontaneous firing, however Dup15q neurons fired at triple the rate of controls. Increased firing rate in Dup15q neurons was confirmed with population calcium imaging. Together these data suggest an additional mechanism of hyperexcitability specific to Dup15q neurons. Pharmacological manipulations suggest that KCNQ2 channels may be impaired in Dup15q neurons. KCNQ2 channels are potassium channels that are modulated by muscarinic receptor activation and act at subthreshold voltages as a brake on repetitive AP firing^42^. Moreover, mutations in KCNQ2 result in benign familial neonatal convulsions, a genetic epilepsy by which children are born with seizures that often attenuate over the first few years of life^42^. Given the physiological role of KCNQ2 and its involvement in seizures, we tested KCNQ2 impairments in Dup15q neurons and found that these cells failed to respond to pharmacological activation or blockade of KCNQ2, suggesting impaired function and/or a reduction of these channels in Dup15q neurons. Interestingly, blockade of KCNQ2 in control neurons resulted in firing rates that were similar to the baseline firing of Dup15q neurons. In line with these data, we found KCNQ2 expression to be significantly diminished in neurons from Dup15q patients as measured by immunostaining/flow cytometry.

Electrophysiological experiments with the KCNQ2 blocker XE991 yield opposite responses in Dup15q and AS neurons, which is interesting in light of the opposite direction of change for *UBE3A* in these patient lines. Though it is less clear whether *UBE3A* is causative in Dup15q, it is believed that it is probably a major role player in Dup15q phenotypes. As our Dup15q patient lines have large duplications encompassing a number of genes, it is unclear whether these phenotypes can be directly linked to increases in *UBE3A*. Recordings from a patient line with paternal duplication to 15q11-q13, in which *UBE3A* is unchanged, but other genes in the region are duplicated, showed responses to both KCNQ2 block and activation that were identical to control neurons. These experiments lend further support to the idea that KCNQ2 may be downstream of *UBE3A* changes, though it remains to be determined how *UBE3A* regulates KCNQ2 expression and function.

It is clear that synaptic dysfunction is an important aspect of the pathophysiology in both AS and Dup15q mouse models, and is also seen in our studies of AS and Dup15q patient-derived neurons. Despite this, the cellular and molecular causes of seizures in these syndromes remain elusive. It has been reported that increased gene dosage of *UBE3A* is directly responsible for a seizure phenotype in a mouse model of Dup15q and that network activity generated by this seizure phenotype interacts with the transynaptic protein Cbln1 in the ventral tegmental area to causes changes in sociability in these mice^14^. Our study identifies three different mechanisms of hyperexcitability: 1) Increased frequency and amplitude of spontaneous synaptic events, 2) impaired synaptic scaling in response to increases in network activity, and 3) increases in spontaneous AP firing as a result of disrupted KCNQ2 channels. Moreover, these mechanisms are distinct from the cellular phenotype of AS-derived neurons and exist in spite of a normal RMP development and only slight developmental delays in AP firing and other intrinsic properties. It is unclear if and how these mechanisms interact to generate seizures in actual brain tissue. Research using animal models has established that impairments in the ratio of excitation to inhibition is an important contributor to ASD and ASD-related pathophysiology, including the high prevalence of seizures in these syndromes^43, 44^, so it is likely that the increases in spontaneous synaptic event frequency and amplitude and impaired scaling are contributors to a similar pathophysiology in Dup15q brains. An important next step requires screening mouse models of Dup15q for similar electrophysiological phenotypes, particularly at very early developmental time points, to link cellular phenotypes to behavior. Furthermore, it remains to be seen whether the synaptic and AP/KCNQ2 phenotypes in Dup15q neurons are causally linked. It is interesting to speculate that disruptions in KCNQ2 function that lead to excessive firing might impair a neurons ability to scale AMPA receptor amplitudes in response to further increases in firing, essentially a ceiling effect. Overall, this study establishes a robust cellular phenotype in human Dup15q neurons with clinically-relevant genetic disruptions that can be used to screen for compounds aimed at reversing hyperexcitability as a foundation for treating seizures in Dup15q patients.

## Acknowledgements

This work was supported by NIH Grants NS078753 and MH094896 (E.S.L.), the Dup15q Alliance (J.J.F.), the Angelman Syndrome Foundation (E.S.L.), and the CT Regenerative Medicine Research Fund (E.S.L., S.J.C., and L.M.L.). We thank Dr. Jon Covault and Dr. Richard Lieberman for providing control cell lines and Dr. Evan Jellison for help with flow cytometry.

