## Supplementary Materials for "Hyperexcitable phenotypes in iPSC-derived neurons from patients with 15q11-q13 duplication syndrome, a genetic form of autism"

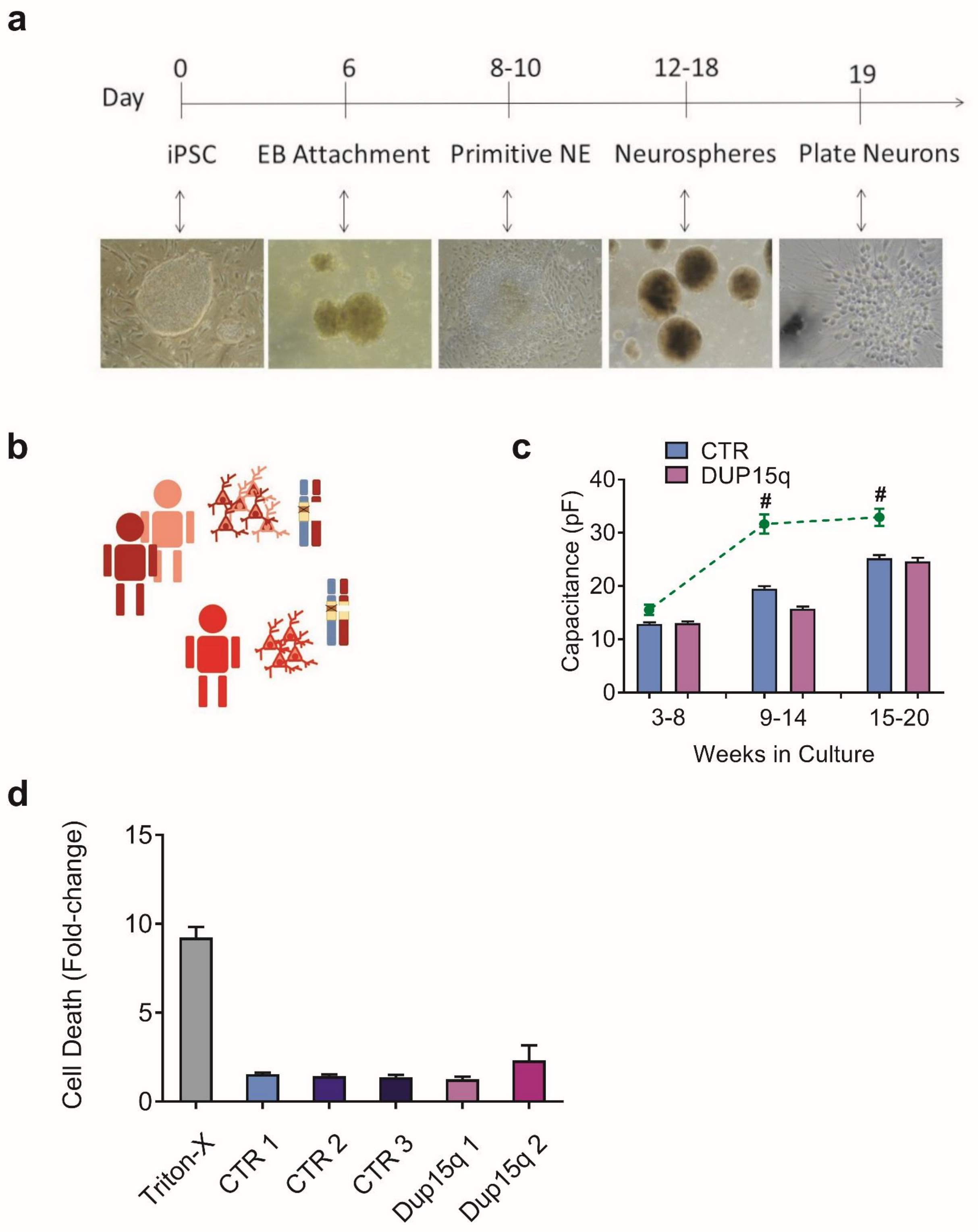
a
Day
0
6
8-10
12-18
19
iPSC
EB Attachment
Primitive NE
Neurospheres
Plate Neurons
b
c
CTR
DUP15q
Capacitance (pF)
40
30
20
10
0
#
#
3-8
9-14
15-20
Weeks in Culture
d
Cell Death (Fold-change)
15
10
5
0
Triton-X
CTR 1
CTR 2
CTR 3
Dup15q 1
Dup15q 2

Figure S1

**Figure S1. Differentiation protocol and additional developmental data. Related to Figure 1.** (**a**) Schematic of neuronal differentiation timeline of induced pluripotent stem cell (iPSC) cultures (**b**) Schematic of Angelman line genetics of the chromosome 15q11-13 genomic locus. (**c**) Group data for cell capacitance for control, Dup15q, and paternal duplication cultures at three time points (n>350 for both control and Dup15q at all time points; n>80 for paternal duplication at all time points). *P<0.001 for significant differences between control and Dup15q, $^{\#}$P<0.001 for significant differences between control and paternal duplication (two-way analysis of variance (ANOVA)). (**d**) Cell death as measured by LDH activity (normalized to cell in plain culture media) for 3 CTR lines (6 samples/line), 2 Dup15q lines (6 samples/line), and control cells treated with Triton-X as a positive control (4 samples).

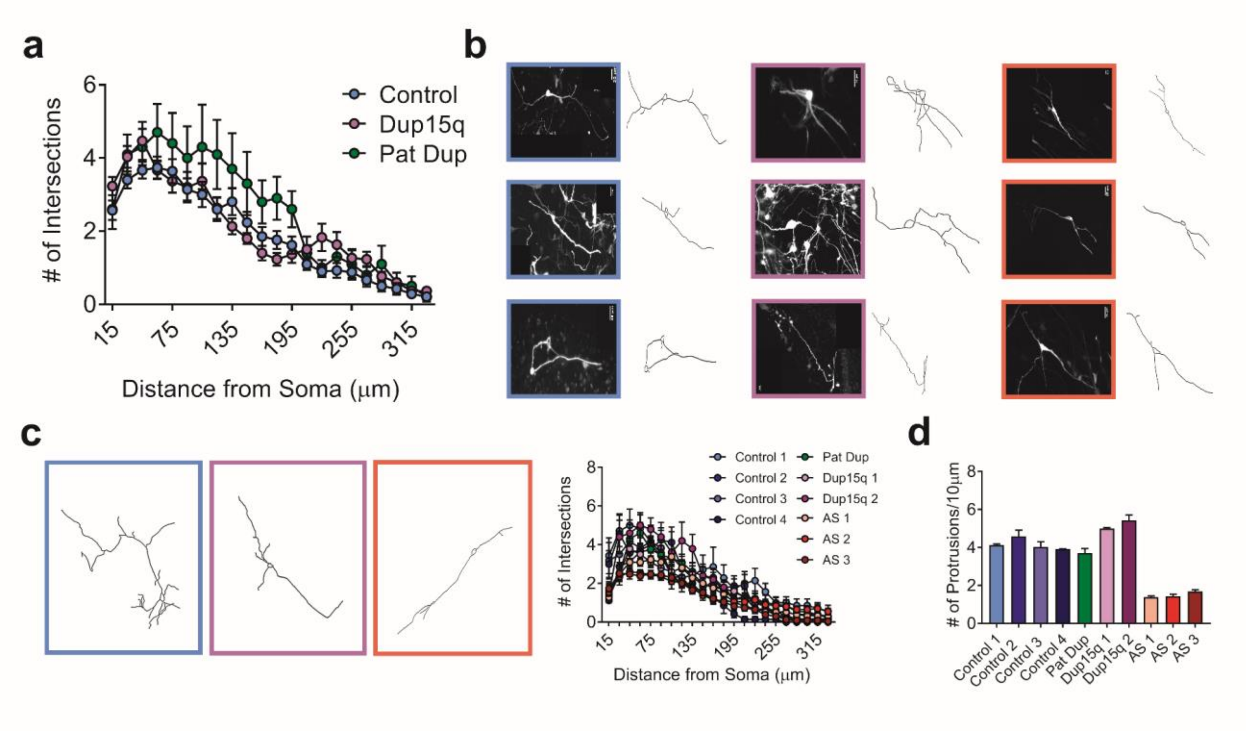


Figure S2

**Figure S2. Additional sholl and protrusion analysis and GluR1/MAP2 colocalization. Related to Figures 2 and 3.** (**a**) Grouped Sholl analysis for neurons (15-20 weeks) from control- (n>25), Dup15q- (n=9), and 15q11-q13 paternal duplication-derived (n=9) cultures. (**b**) Additional example images and respective Neuron Tracer images of transfected neurons used for analysis from control (left; blue), Dup15q (purple; middle), and Angelman (red; right). Scale bars: 25µm. (**c**) Left: Neuron tracer images of example neurons showed in main Figure 2e. Right: sholl analysis for individual lines used for group data depicted in main Figure 2e (n>8 for each data point). (**d**) Analysis for the number of protrusions (see Methods for details) for individual lines used for group data depicted in main Figure 2f (n>8 for each bar).

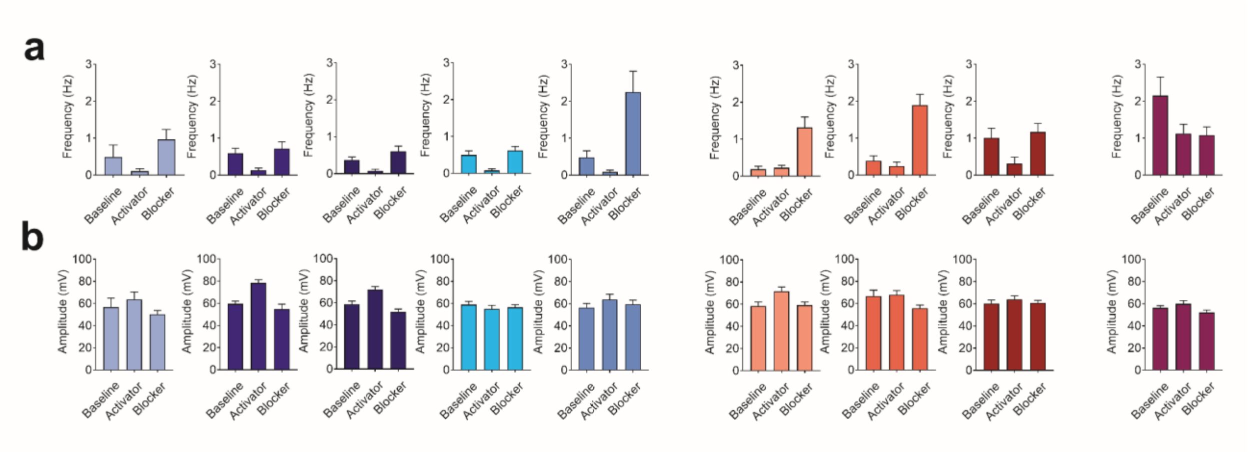


Figure S3

**Figure S3. Additional data for KCNQ activation, blockade, and immunocytochemistry. Related to Figure 6.** (**a**,**b**) Frequency (**a**) and amplitude (**b**) of spontaneous action potential firing for additional control (left), Angelman (middle) and Dup15q (right) lines at baseline or in the presence of either pharmalogical blocker or activator of KCNQ2 channels (each bar represents data from 1-2 coverslips and n=15-30 neurons).
